# Temporal stability of fMRI in medetomidine-anesthetized rats

**DOI:** 10.1101/667659

**Authors:** Nikoloz Sirmpilatze, Jürgen Baudewig, Susann Boretius

## Abstract

Medetomidine has become a popular choice for anesthetizing rats during long-lasting sessions of blood-oxygen-level dependent (BOLD) functional magnetic resonance imaging (fMRI). Despite this, it has not yet been established how commonly reported fMRI readouts evolve over several hours of medetomidine anesthesia and how they are affected by the precise timing, dose, and route of administration. We used four different protocols of medetomidine administration to anesthetize rats for up to six hours and repeatedly evaluated somatosensory stimulus-evoked BOLD responses and resting state functional connectivity throughout. We found that the temporal evolution of fMRI readouts strongly depended on the method of administration. Protocols that combined an initial medetomidine bolus (0.05 mg/kg) together with a subsequent continuous infusion (0.1 mg/kg/h) led to temporally stable measures of stimulus-evoked activity and functional connectivity. However, when the bolus was omitted, or the dose of medetomidine lowered, the measures attenuated in a time-dependent manner. We conclude that medetomidine can sustain consistent fMRI readouts for up to six hours of anesthesia, but only with an appropriate administration protocol. This factor should be considered for the design and interpretation of future preclinical fMRI studies in rats.

## Introduction

Functional magnetic resonance imaging (fMRI), relying on blood-oxygen-level dependent (BOLD) contrast^1^, is being widely used for the non-invasive mapping of human brain function. Classically, fMRI has focused on the brain’s response to a task or stimulus, but more recent task-free (resting state fMRI) approaches have explored spontaneous low frequency fluctuations in the BOLD signal^2,3^, and their role in the functional connectivity of healthy and diseased brains4. Since the advent of dedicated high-field MR systems, fMRI applications have expanded to experimental animals, especially rodents^5^. This is a promising development for translational preclinical research, since the same technique can be applied to both human patients and animal models. Additionally, the expanding capacity of small animal fMRI to be combined with genetic, pharmacological and surgical manipulations, as well as with electrophysiological and optical recordings, allows the exploration of increasingly complex neuroscientific questions^6–9^.

Nevertheless, small animal fMRI poses a methodological challenge: it necessitates the subject’s immobility for long imaging times. Animals can be restrained and habituated to head fixation and MR scanner noise, but this is laborious for the researcher, and often stressful for the animal^10–12^. Thus, ethical and practical considerations mandate the use of anesthesia in the majority of small animal fMRI studies. Unfortunately, anesthetics confound fMRI measurements in multiple ways: they may alter neural activity, affect systemic cardiorespiratory physiology, and interfere with cerebral vasculature and neurovascular coupling—the very mechanism giving rise to the BOLD contrast^13,14^. Therefore, there is a need for an anesthetic protocol that ideally provides sufficient, long-lasting sedation, while maintaining neural activity and neurovascular coupling. In pursuit of the above properties, researchers have tried multiple anesthetic agents, including α-chloralose, medetomidine, isoflurane, propofol, urethane, and ketamine-xylazine. These agents have varying effects on the neurovascular system, with each of them presenting a unique set of benefits and drawbacks for fMRI applications^12–18^. This has led to a substantial diversity in anesthetic protocols used for rodent fMRI, compromising the comparability and reproducibility of results.

Of the above anesthetics, the sedative agent medetomidine—a highly selective α_2_-adrenergic agonist—holds perhaps the most promise for becoming a routine choice for fMRI applications in rats. Medetomidine comes as an equal mixture of two enantiomers, with the dextro-isomer, dexmedetomidine, being the active component^19,20^. Dexmedetomidine decreases the activity of noradrenergic neurons in the locus coeruleus, producing a state that mimics non-REM sleep^21^. Conveniently, the drug’s effects are reversible by a specific α_2_-adrenergic antagonist— atipamezole^19,20^. A protocol based on the continuous infusion of medetomidine, first introduced by Weber et al. in 2005^22^, presents several advantages for fMRI studies: it sedates rats for several hours, leads to robust stimulus-evoked BOLD responses, allows for easy subcutaneous (SC) administration, avoids the need for intubation, and can be used for longitudinal studies with multiple fMRI sessions^23^. Other research groups have since confirmed the benefits of medetomidine infusion and expanded its usage to resting state fMRI^24–27^. Medetomidine anesthesia is currently an established practice for rat fMRI, with at least 40 published articles reporting the use of the original medetomidine protocol, or variations of it (Supplementary Table S1). Its use is expected to rise, owing to the practical advantages and to the increasing availability of techniques that can be combined with rat fMRI.

However, there are several concerns revolving around the duration and stability of medetomidine anesthesia. Medetomidine administration is always preceded by an inhalable gas anesthetic, usually isoflurane, which is used for anesthesia induction and animal preparation. Isoflurane alters brain metabolism^28^, strongly suppresses stimulus-evoked hemodynamic responses^29–31^, and may introduce widespread correlations in functional connectivity metrics^12,27,32^; the possibility of these effects lingering long after isoflurane discontinuation cannot be excluded^33^. Then there is the issue of medetomidine itself, which is usually given in two steps: first as a bolus loading dose and then as a continuous infusion. Due to the drug’s strong α_2_-adrenergic effects^20^, these actions are expected to time-dependently alter hemodynamic parameters, which could in turn affect BOLD-based readouts. Another important concern is the restricted duration of anesthesia, with animals reported to spontaneously wake up despite the continuous infusion of medetomidine. Tolerance to the drug’s sedative effects has been blamed for this, with researchers proposing to counter it by stepping up the infusion rate^25^. All the above issues are further compounded by the lack of consensus regarding the exact administration scheme; protocols vary in administration route and dose, while some researchers choose to omit the bolus (see Supplementary Table S1).

These concerns imply that commonly reported fMRI readouts, namely stimulus-evoked BOLD responses and resting state functional connectivity (RSFC), might not be stable over long-lasting imaging sessions. It is unclear when a steady state is reached by these readouts, for how long it is maintained, and how it is affected by various medetomidine administration choices. An answer to these questions would enable researchers to design rat fMRI experiments in a way that maximizes the duration of the steady state. This would increase the available experimental time, decrease variance, and ultimately reduce the total number of required animals, all the while promoting comparability among studies.

To achieve the above goals, we examined how stimulus-evoked BOLD responses and RSFC evolve over time during medetomidine anesthesia. We tested four different protocols for medetomidine administration (Table 1, Fig. 1a) on two separate sessions: firstly on a laboratory bench and secondly inside a small animal MR-system. During the latter session, we performed multiple repeated fMRI measurements (runs), with consecutive runs being alternated between somatosensory stimulus-evoked fMRI with electrical forepaw stimulation (EFS-fMRI) and resting state fMRI (RS-fMRI) (see Fig. 1b-c). The acquired runs, 283 EFS-fMRI and 295 RS-fMRI in total, spanned a period of 0.5 – 6 hours relative to the start of medetomidine administration. For each medetomidine protocol, we report the achieved duration of anesthesia and the following measures across time: heart and respiratory rates (HR and RR); localization and amplitude of stimulus-evoked responses; strength and structure of RSFC. Based on our findings, we make recommendations regarding the administration protocol of medetomidine and the timing of fMRI experiments within the protocol.

**Table 1.**
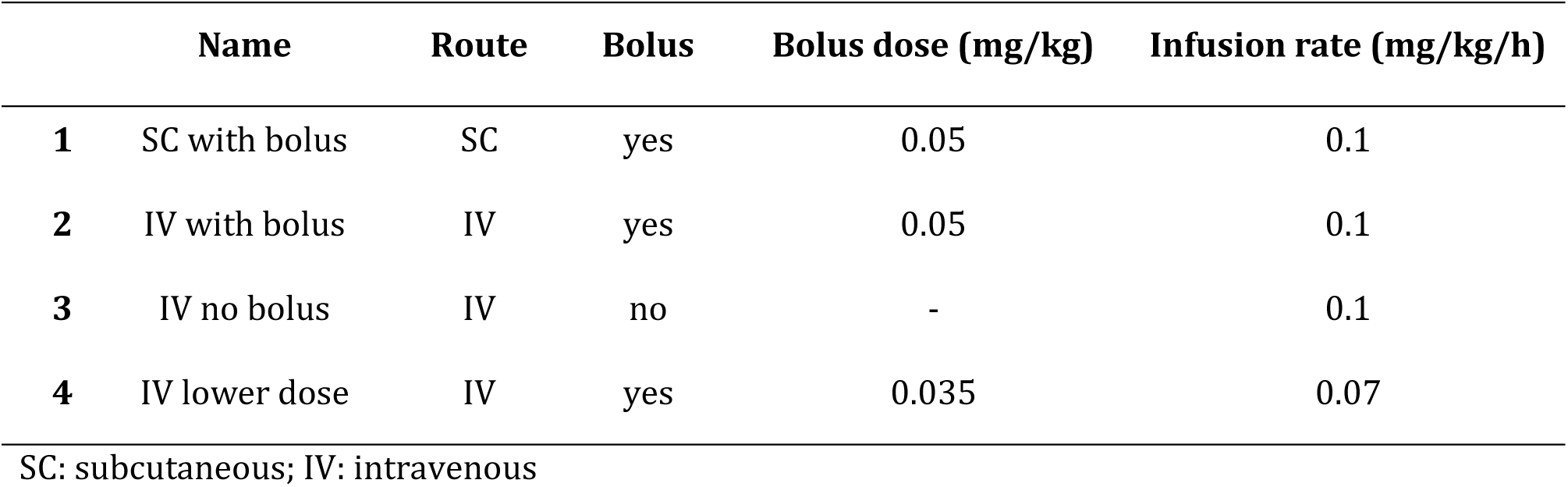
Medetomidine administration protocols used in this study.

|  | Name | Route | Bolus | Bolus dose (mg/kg) | Infusion rate (mg/kg/h) |
| --- | --- | --- | --- | --- | --- |
| 1 | SC with bolus | SC | yes | 0.05 | 0.1 |
| 2 | IV with bolus | IV | yes | 0.05 | 0.1 |
| 3 | IV no bolus | IV | no | - | 0.1 |
| 4 | IV lower dose | IV | yes | 0.035 | 0.07 |

SC: subcutaneous; IV: intravenous

**Fig. 1.**
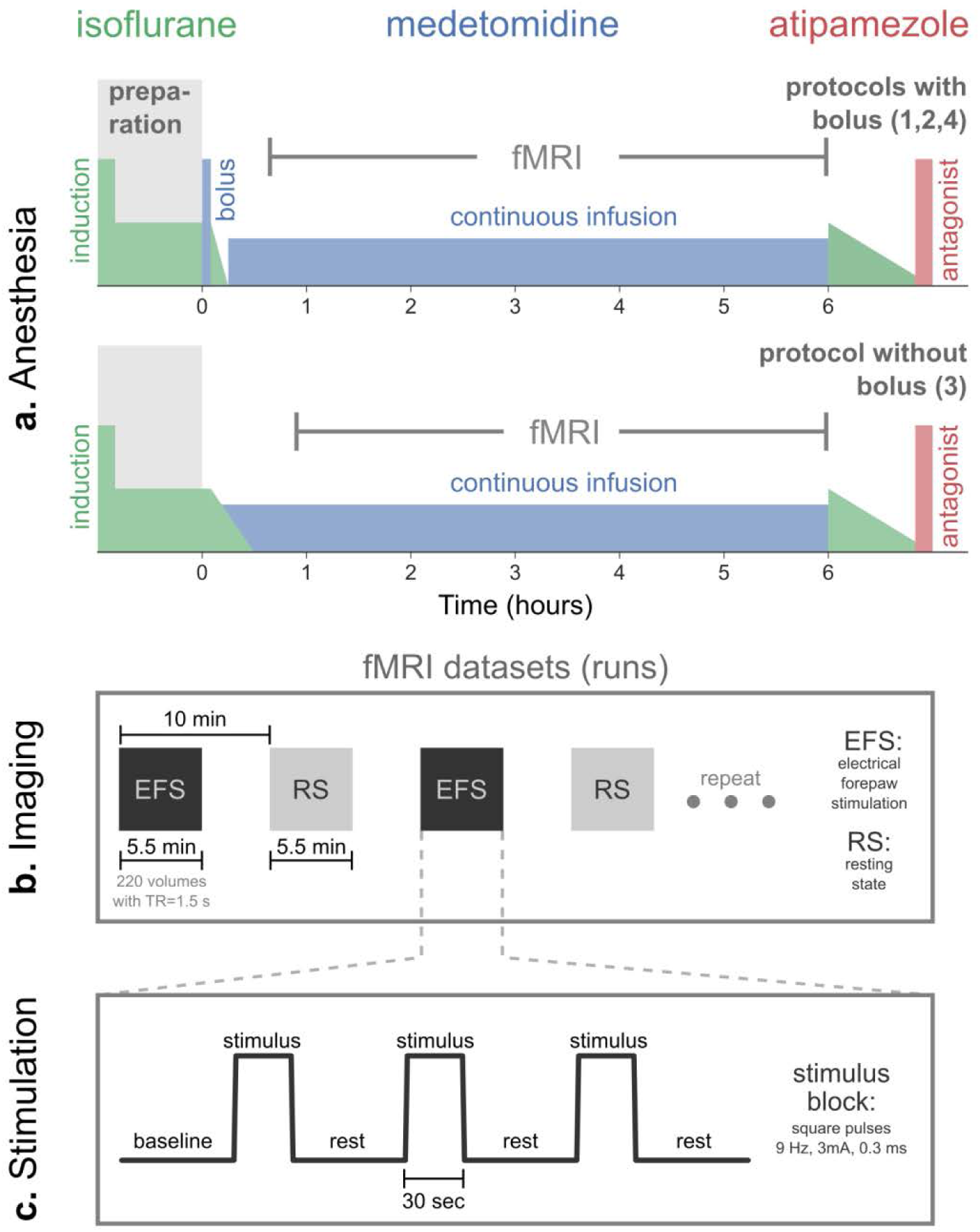
Anesthetic protocols and fMRI acquisition. **(a)** The general outline of the applied anesthetic protocols. For all four protocols (see Table 1), isoflurane was used to induce unconsciousness (5%) and during animal preparation (2-3%). For protocols 1, 2, and 4, a bolus of medetomidine was given after the preparation phase, followed by a gradual reduction of isoflurane and its eventual discontinuation 10 min later; continuous infusion of medetomidine commenced 15 min after the bolus. The bolus was omitted for protocol 3, with continuous infusion starting directly after preparation, and isoflurane being gradually reduced to zero over the course of 20 – 25 min. Anesthesia was maintained for a maximum of six hours since the start of medetomidine administration (time = 0). In the end, animals were provided with 2% isoflurane and freed from all equipment. Atipamezole was injected SC to antagonize medetomidine effects and to facilitate a smooth recovery. **(b)** Multiple fMRI runs were acquired per anesthesia session, with consecutive runs being alternated between somatosensory fMRI with electrical forepaw stimulation (EFS-fMRI) and resting state fMRI (RS-fMRI) with no stimulus. The stimulation paradigm applied during EFS-fMRI runs is shown in **(c)**.

## Results

### Anesthesia duration and physiology

Out of all 48 anesthesia sessions, 27 lasted for the full six hours, while 21 ended with a spontaneous wake-up (spontaneous movement for bench sessions; rapid rise in RR for fMRI sessions). These wake-up incidents occurred across session types (8/24 bench; 13/24 fMRI) and medetomidine protocols (4, 6, 7, and 4 out of 12, for protocols 1 – 4 respectively). However, only a few of those occurred early, with 35/48 sessions (72.9%) exceeding five hours in duration (Fig. 2, left). The HR and RR followed similar temporal trends across all four medetomidine protocols (Fig. 2, right). HR decreased rapidly after the introduction of medetomidine, dropping by approximately 50% within the first hour of anesthesia. After that it showed only a slight tendency to gradually recover over time. The RR also underwent rapid changes in the first hour: it started at 40-70 bpm under isoflurane, decreased in response to medetomidine introduction, and gradually recovered following the discontinuation of isoflurane. As was the case with HR, it remained mostly stable after the first hour.

**Fig. 2.**
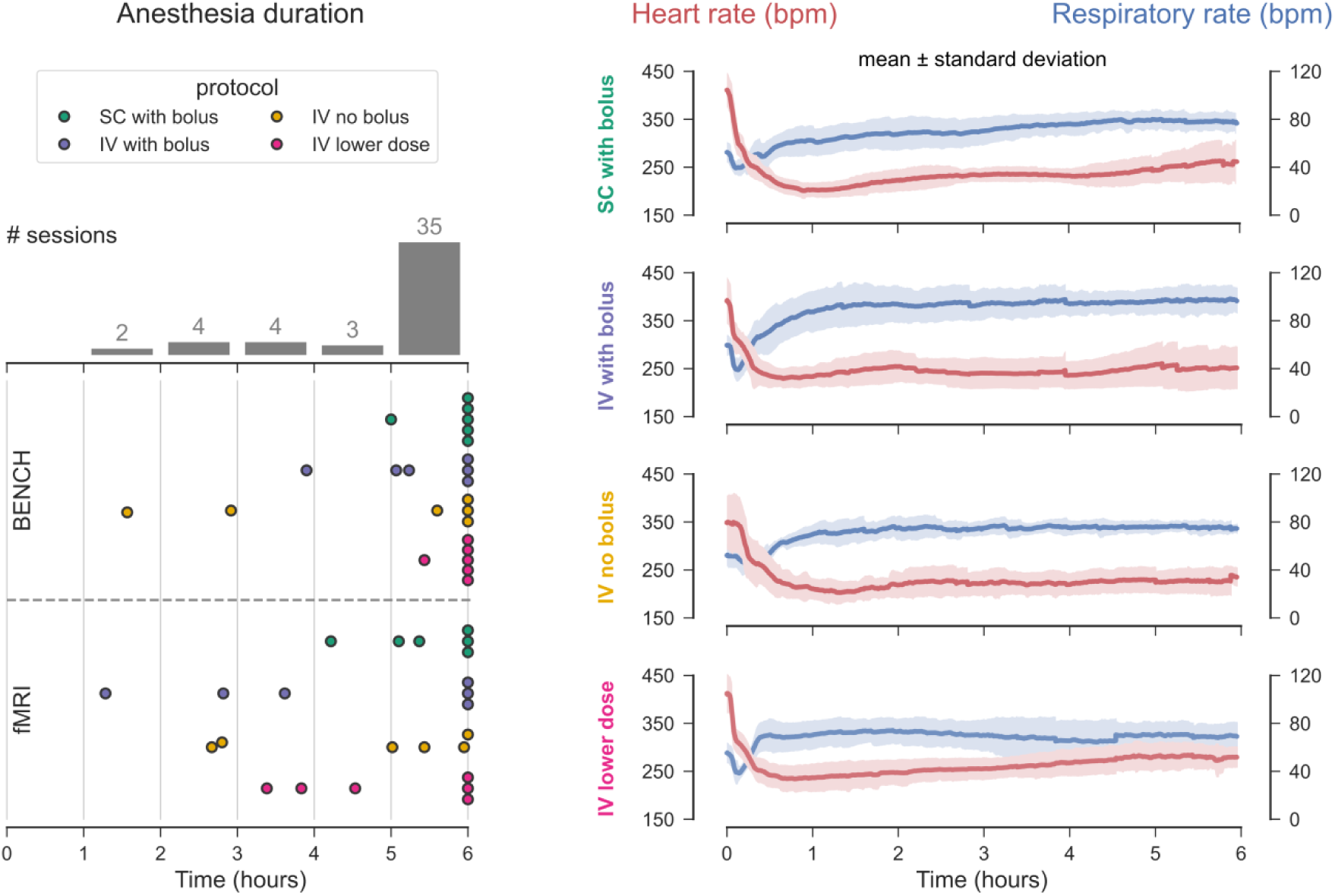
Anesthesia duration and physiology. **(a)** Anesthesia duration is plotted (dots) separately for bench and fMRI sessions, across all four medetomidine protocols. The bars represent a histogram of anesthesia durations, with all 48 anesthesia sessions binned into one-hour intervals. **(b)** Heart and respiratory rates (in beats/breaths per minute—bpm), pooled from both bench and fMRI sessions of each medetomidine protocol, are plotted as an across-session mean (solid line) ± s.d. (shaded area).

### Areas activated by the stimulus

For each EFS-fMRI run we identified the active areas using a first-level general linear model analysis. The thresholded statistical maps (cluster threshold, z > 3.1, p = 0.05) were binarized (1 for active voxels, 0 elsewhere) and averaged across all EFS-fMRI runs to produce an activation probability map. The probability map revealed a consistently active cluster, anatomically corresponding to the left (contralateral to the stimulus) forelimb region of the primary somatosensory cortex—abbreviated as S1FL (Fig. 3a left). This cluster’s center was active in 85.16% of all EFS-fMRI runs, whereas no other area was active in more than 7% of runs. To ensure that there was no systematic shift in the location of the active S1FL cluster, the 283 first-level activation maps were split into 12 groups according to the applied medetomidine protocol and the time since the start of medetomidine administration (early: 0 – 2 h; middle: 2 – 4 h; late: 4 – 6 h). Examination of activation probability maps from all groups (Fig. 3a, right) verified that the active cluster’s location remained stable across time and medetomidine protocols.

**Fig. 3.**
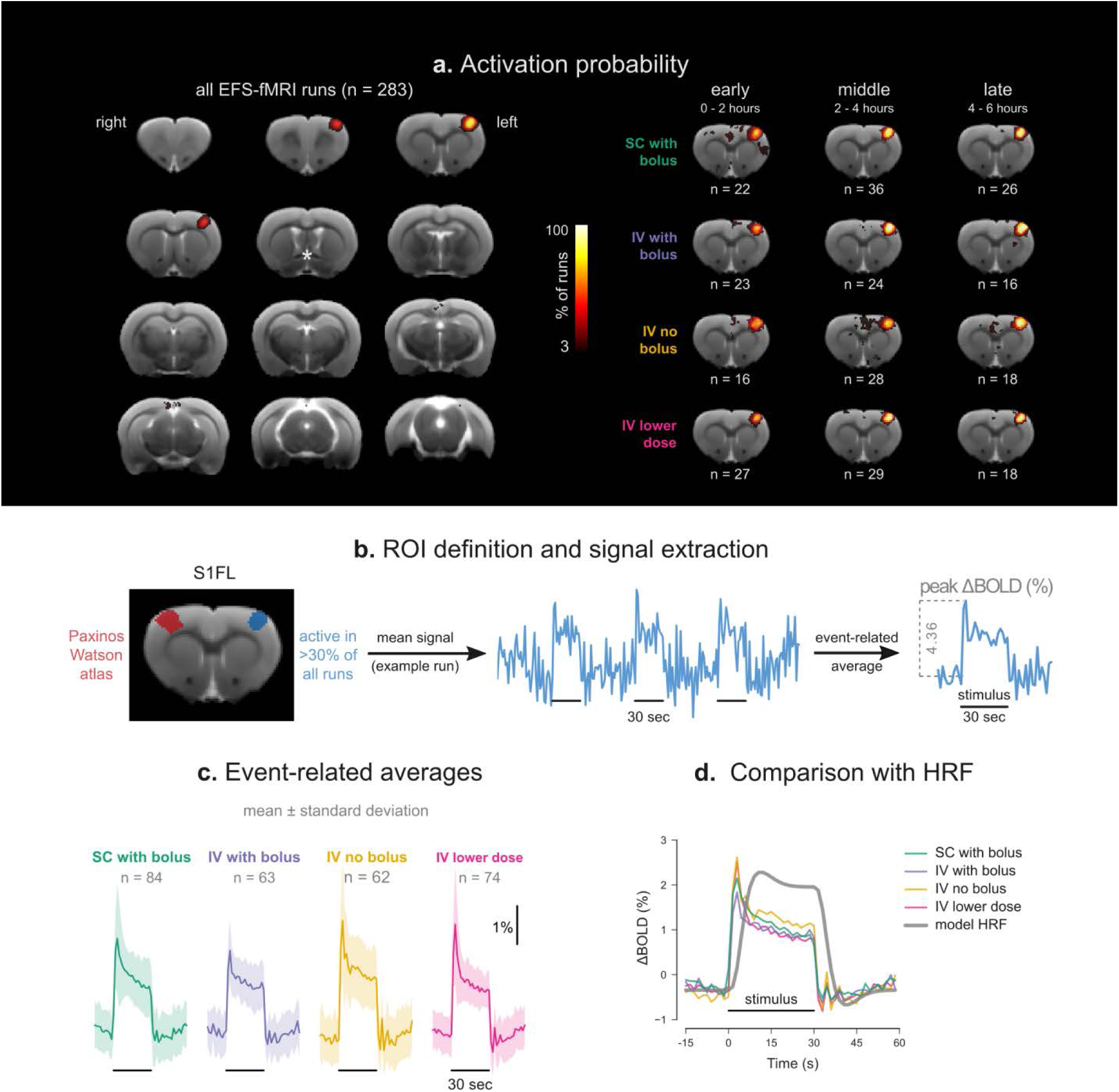
Areas activated by electrical forepaw stimulation (EFS). **(a)** An activation probability map produced by pooling significantly active clusters across all 283 EFS-fMRI runs (left). On the right, the EFS-fMRI runs are grouped according to the applied medetomidine protocol and to the time since the start of medetomidine administration (early: 0 – 2 h; middle: 2 – 4 h; late: 4 – 6 h), to produce separate activation probability maps for each group. All maps are thresholded at 3% and overlaid on a T2-weighted structural study template. The asterisk marks the crossing of the anterior commissure (AC, −0.36 mm relative to the bregma, according to the Paxinos-Watson rat brain atlas). The rest of the slices shown on the left are taken at 1 mm intervals from the AC slice. On the right, only the slice containing the peak activation, 2 mm rostral to AC, is shown for each group. **(b)** The only consistently active cluster across all runs corresponds to the forelimb region of the left primary somatosensory cortex (S1FL). The location of this cluster is shown alongside the anatomical delineation of the same area from the Paxinos-Watson rat brain atlas. The functionally defined S1FL (area active in > 30% of all EFS-fMRI runs) is set as a region-of-interest (ROI) for the extraction of BOLD signal time courses. Such a time course is shown for one example EFS-fMRI run, with the stimulation blocks marked by horizontal lines. Averaging the three stimulation blocks results in an event-related average, from which the peak % signal change (peak ΔBOLD) can be extracted. Event-related average responses (mean ± s.d.) are plotted for all four medetomidine protocols **(c)** and compared to the model response **(d)**. The model is generated by convolving the stimulus paradigm (1 during EFS, 0 otherwise) with a double-gamma hemodynamic response function (using the default parameters of FEAT version 6.00, part of FSL); it is shown here rescaled to the y-axis range of the event-related averages, to aid visual comparison.

### Shape and strength of stimulus-evoked responses

The portion of the S1FL that was significantly active in at least 30% of all 283 EFS-fMRI runs was taken as a functionally-defined region-of-interest (ROI). For each EFS-fMRI run, this ROI’s mean BOLD time course was extracted, normalized to the pre-stimulus baseline, and averaged across stimulation blocks to produce an event-related average. This was used to extract the peak % signal change (peak ΔBOLD), as a measure of stimulus-evoked BOLD response strength (Fig. 3b). The examination of event-related averages revealed that the shape of S1FL BOLD responses was very similar across the four medetomidine protocols (Fig. 3c). The signal exhibited a sharp peak after stimulus onset, followed by a plateau lasting till the end of the 30 s stimulation. The time-to-peak was found to be consistently about 3 s for all protocols—shorter than what is assumed by the default hemodynamic response functions of most fMRI analysis packages (Fig. 3d).

The peak ΔBOLD exhibited varying temporal trends depending on the medetomidine protocol (Fig. 4a-b). Protocol 2 (IV with bolus) led to the most stable BOLD responses, with mean peak ΔBOLD of 2.4%, independent of time (p = 0.61, Likelihood Ratio Test). Omitting the bolus (protocol 3, IV no bolus), or lowering the medetomidine dose (protocol 4, IV lower dose), led to stronger early responses at around 4 – 4.5%, but these were not sustainable over time: peak ΔBOLD decayed with negative slopes of 0.43%/hour (p < 0.001) and 0.47%/hour (p < 0.001) respectively. Protocol 1 (SC with bolus) also exhibited a decreasing temporal trend, but with a smaller slope (−0.24%/hour, p = 0.02), and with the effect being driven by four exceptionally strong early BOLD responses in two of the rats. In general, and despite their differing temporal trends, all four protocols converged to a mean peak ΔBOLD of around 2.5% after two hours of medetomidine anesthesia (Fig. 4c).

**Fig. 4.**
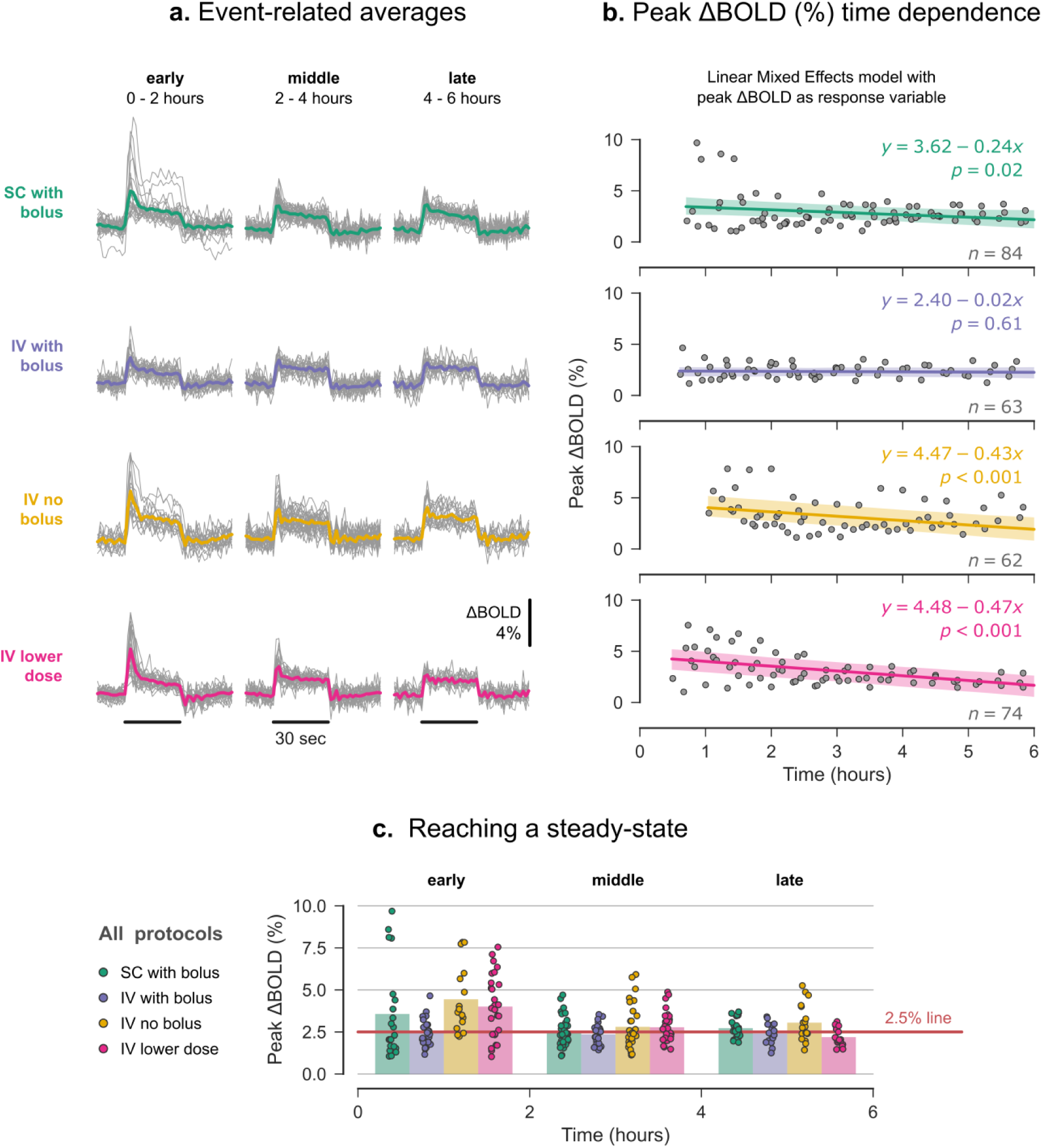
Temporal stability of stimulus-evoked responses. **(a)** Event-related averages produced by averaging the three electrical forepaw stimulation (EFS) blocks of each fMRI run. Single run traces are grouped according to the applied medetomidine protocol and the time since the start of medetomidine administration (early: 0 – 2 h; middle: 2 – 4 h; late: 4 – 6 h); each group’s mean event-related average is plotted as a thicker trace; EFS duration is represented by horizontal lines. The peak stimulus-evoked signal change (peak ΔBOLD) is extracted from the event-related averages and plotted in **(c)**; dots represent single run values, while bars show the group mean. The same single-run peak ΔBOLD values are plotted in **(b)** against the exact time of data acquisition. Solid lines correspond to the fit of a Linear Mixed Effects model (equation shown), with peak ΔBOLD as a response variable, time as a fixed effect, and individual rat intercepts as random effects; shaded areas represent the 95% confidence intervals of the fit. To test for time dependence, the constructed models were compared to corresponding models lacking the time effect, using a Likelihood Ratio Test; the resulting p-values are shown.

### Resting state functional connectivity

RSFC was probed by examining the pair-wise correlations between the BOLD time courses of 28 anatomically defined ROIs (Fig. 5a). Examination of the pair-wise correlation matrices (Fig. 5b), and of their network representations (Fig. 5c), showed that the hierarchical structure of the network, i.e. the strength of individual connections relative to each other, was consistent over time for all medetomidine protocols. The network’s global RSFC—the mean correlation (Fisher’s Z-score) across all unique ROI pairs—was computed for all RS-fMRI runs, and tested for time dependence (Fig. 5d). Bolus-based medetomidine administration (protocols 1, 2 and 4) provided stable global RSFC throughout the six hours of anesthesia: Z-scores remained at 0.57 ± 0.27 (mean ± s.d.), 0.44 ± 0.17, and 0.34 ± 0.12 respectively, and exhibited no significant temporal trends (p = 0.80 for protocol 1; p = 0.17 for protocol 2; p = 0.09 for protocol 4). When the bolus was omitted (protocol 3, IV no bolus), global RSFC was stronger during the early period (0.85 ± 0.30), but decreased time-dependently with a slope of −0.07/hour (p = 0.02).

**Fig. 5.**
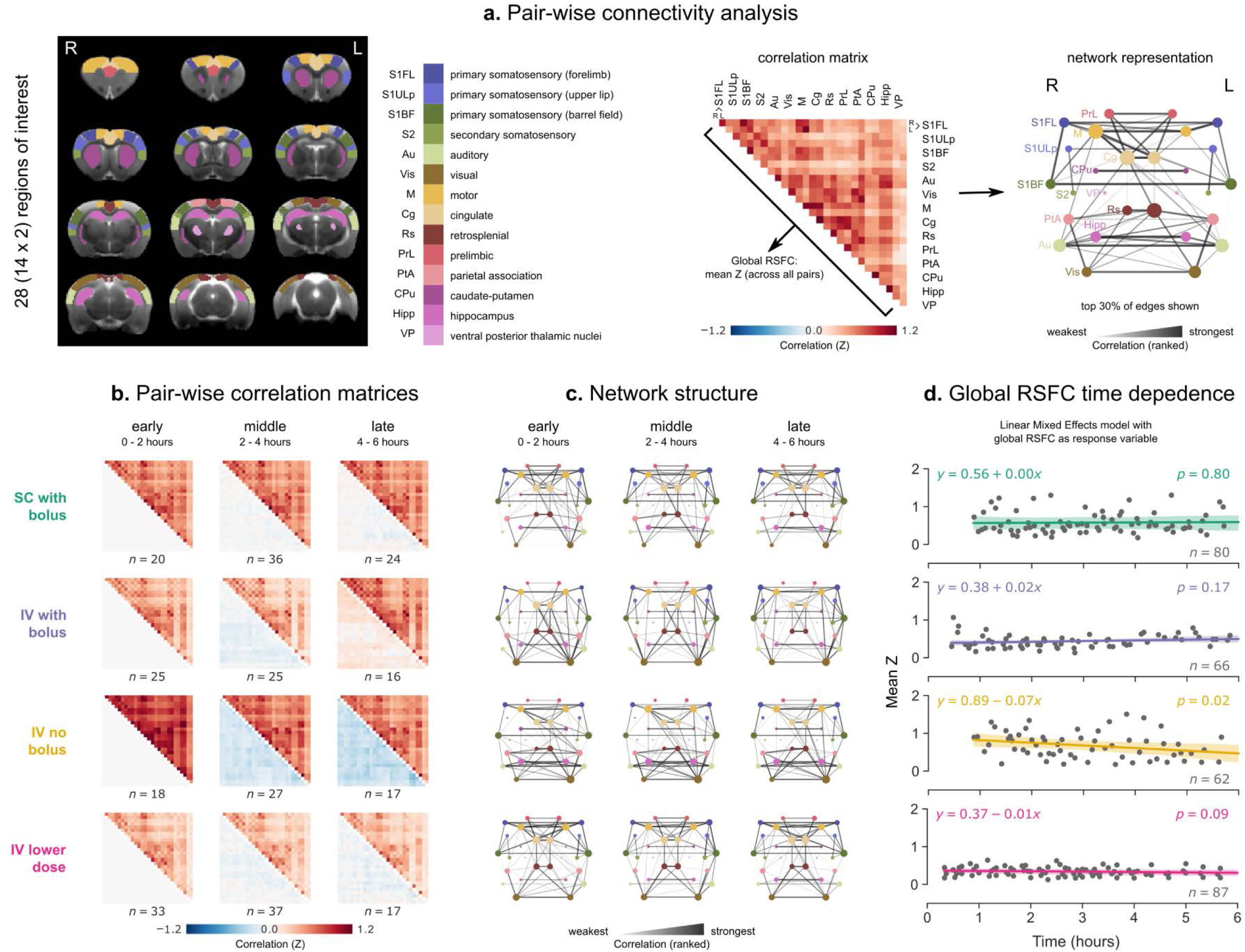
Resting state functional connectivity (RSFC). **(a)** To calculate pair-wise RSFC, 28 regions-of-interest (ROIs) were defined based on the Paxinos-Watson rat brain atlas—14 on each hemisphere. Pearson’s correlations were calculated between the BOLD time courses of all unique ROI pairs and transformed into Fisher’s Z-scores. A pair-wise correlation matrix is shown for one example RS-fMRI run; the mean correlation (Z-score) across all ROI pairs constitutes the global RSFC. The matrix is also represented as a weighted network graph, with the ROIs as nodes and their pairs as edges. For visualization clarity, only the strongest 30% of edges are shown; edge thickness and opacity scale linearly with the relative rank of the correlation value (the highest Z-score corresponds to the thickest edge); node radius scales with the weighted degree (weighted sum of edges passing through the node). **(b)** The 295 RS-fMRI runs are split into 12 groups according to the applied medetomidine protocol and the time since the start of medetomidine administration (early: 0 – 2 h; middle: 2 – 4 h; late: 4 – 6 h). For each group, the upper triangular matrix represents the mean (across runs) pair-wise correlation, while the lower triangular matrix shows the change in correlation compared to each protocol’s early period. **(c)** The mean correlation matrices (upper triangles) are also visualized as network graphs, similarly to the example graph above. **(d)** Global RSFC values (dots) are plotted against the time of the RS-fMRI run acquisition. Solid lines correspond to the fit of a Linear Mixed Effects model (equation shown), with global RSFC as a response variable, time as a fixed effect, and individual rat intercepts as random effects; shaded areas represent the 95% confidence intervals of the fit. To test for time dependence, the constructed model was compared to a corresponding model lacking the time effect, using a Likelihood Ratio Test; the resulting p-values are shown.

## Discussion

In the present study we evaluated the capacity of medetomidine—administered through four different protocols—to anesthetize rats for up to six hours and sustain temporally stable fMRI measures of stimulus-evoked activity and functional connectivity. We found that anesthesia duration exceeded five hours in most sessions (35/48 or 72.9%; Fig. 2, left), heart and respiratory rates were mostly stable after the first hour of anesthesia (Fig. 2, right), and the majority of performed EFS-fMRI runs (241/283 or 85.16%) led to a significant and selective activation of the expected cortical area (Fig. 3a). However, the temporal stability of both stimulus-evoked activity and functional connectivity was observed to strongly depend on the protocol of medetomidine administration (Fig. 4 and 5).

Limited anesthesia duration has been referred to as a drawback of medetomidine protocols^34^, even though the exact duration is not reported in most papers. According to a study that has addressed this issue, rats anesthetized by constant IV infusion woke up spontaneously 3.5-4 hours into the experiment^25^. However, the rats in that study were tracheotomized and mechanically ventilated, while most rat fMRI experiments—including ours—are conducted in freely breathing animals. The anesthesia durations achieved in our experiments should satisfy the needs of most researchers. That said, the risk of spontaneous wake-ups must be taken into consideration. In our study, these wake-ups occurred more often in fMRI than in bench sessions (13/24 versus 8/24)—a possible effect of the scanner’s acoustic noise, and were typically preceded by an increase in RR and arrhythmic breathing. If early wake-ups have to be avoided, other administration practices may be warranted, such as stepping up the infusion rate^25^, or adding a constant low dose of isoflurane throughout the imaging session^35^. The latter strategy has gained popularity in recent years, on the basis that it provides near-normal physiological conditions^35^ and protects against epileptic seizures—which have been reported in animals with medetomidine-only anesthesia^36^. However, this might require mechanical ventilation, since even low doses of isoflurane, when combined with medetomidine, have been shown to suppress the amplitude of stimulus-evoked BOLD responses in spontaneously breathing rats^37^.

The recorded cardiorespiratory parameters (HR and RR) underwent rapid changes in the first hour of anesthesia, but stabilized thereafter (Fig. 2, right). The observed changes agree with medetomidine pharmacology and with previous rat fMRI studies^20,23,38^. The decrease in HR immediately after medetomidine injection can be attributed to its well-described α^2^-adrenergic effects. Since the respiratory effects of medetomidine alone are considered to be minor, the observed initial drop in RR was probably caused by medetomidine enhancing the potency of isoflurane—and thus amplifying isoflurane-induced respiratory depression^20^. This interpretation is supported by the gradual recovery of RR following the discontinuation of isoflurane.

The EFS-evoked activation of the contralateral S1FL (Fig. 3) is described by numerous other rat fMRI studies using the same stimulus^23–25^. For a subset of trials, these studies have also reported activations in the secondary somatosensory area and in sensory thalamic nuclei. If we increase statistical power by grouping multiple fMRI runs in a second-level analysis, we also find the same areas being responsive to EFS (see Supplementary Fig. S2). The shape of the EFS-evoked BOLD response in the S1FL (sharp peak at about 3 s after the onset of stimulation, followed by a plateau sustained till the end of EFS period; see Fig. 3c) is consistent with previous rat fMRI studies^24,39^, but differs considerably from what would be expected based on the standard human double-gamma hemodynamic response function^40^. Specifically, the response in medetomidine-anesthetized rats exhibits faster kinetics (Fig. 3d). The reasons for that could be diverse, including the different species, the use of anesthesia, the higher magnetic field strength utilized in rodent studies, or a combination of these factors. Nevertheless, since rat fMRI with medetomidine is becoming such a common practice (see Supplementary Table S1), future studies should focus on determining the optimal response function for this paradigm.

The temporal evolution of peak stimulus-evoked responses exhibited an interesting dependence on the medetomidine administrations protocol (Fig. 4): the peak ΔBOLD differed significantly among protocols in the early two hours of anesthesia, before eventually converging to the same value. Specifically, combining a bolus of 0.05 mg/kg with a subsequent continuous infusion of 0.1 mg/kg/h—the most widely used medetomidine dosage in rat fMRI studies—led to temporally stable responses, especially when administered via the IV route. Deviating from the above dosage, either by omitting the bolus (protocol 3) or by downscaling both bolus and infusion doses (protocol 4), led to significantly stronger early responses. Omitting the bolus also had a similar effect on global RSFC: the overall pair-wise correlation strength decreased over time. This time-dependent attenuation of BOLD readouts in sessions that lacked the bolus could reflect a negative relationship between the strength of these readouts and the concentration of medetomidine. Drug levels in the central nervous system might take hours to stabilize under the continuous infusion regime, given that the pharmacokinetics of medetomidine is characterized by a long terminal half-life of about 57 min, and a hysteresis between plasma and cerebrospinal fluid concentrations^38,41^. Bolus administration likely mediates a faster wash-in of the drug and an earlier establishment of the steady state. For this interpretation, we need to accept that medetomidine dose-dependently suppresses stimulus-evoked BOLD responses and RSFC, at least up to a certain level. Previous studies have in fact found such a dose-dependency for RSFC, but not for stimulus-evoked responses^39,42^. That said, the relevant experiments were restricted to a higher infusion rate range of 0.1 – 0.3 mg/kg/h (always preceded by a bolus), within which any effects on stimulus-evoked responses could have been saturated.

At this point we would like to emphasize several limitations of the present study. Importantly, we have not measured blood concentrations of medetomidine, and thus we cannot directly verify our interpretation of the dose-dependent effects. Moreover, we have no way of dissecting the vascular and neuronal contributions to the observed effects on BOLD readouts: we lack electrophysiological recordings of neural activity and important measures of vascular physiology, such as cerebral blood flow and arterial blood gas concentrations. We also lack fMRI data for very early time points: the earliest fMRI runs were acquired 30 – 50 minutes after the onset of medetomidine administration, due to the time spent for animal positioning, structural image acquisition and shimming, and the required overlap between medetomidine administration and isoflurane anesthesia. This delay was most pronounced for protocol 3, during which the said overlap was necessarily prolonged.

Despite these limitations, our study does provide an empirical answer to the question of how BOLD readouts evolve over long-lasting medetomidine anesthesia sessions, allowing us to recommend best practices for maximizing the duration of the steady state. Whenever temporal stability is deemed crucial, a bolus administration of medetomidine and a dosage according to the originally published protocol^23^ are strongly recommended. Both IV and SC routes can be considered: IV administration leads to less variance in the initial two hours, but SC is easier to implement. Although we lack fMRI data from the first hour of anesthesia, we would advise against performing functional imaging within this period, considering the rapid changes in cardio-respiratory physiology. In conclusion, the recommended dosage (protocols 1 and 2) is clearly sufficient to provide consistent measures of stimulus-evoked activity and functional connectivity in the time period from one to six hours since the bolus. We do not claim that this administration protocol is necessarily the best choice for rat fMRI, since our study did not include all existing (see Supplementary Table S1) or possible administration protocols. It is however reasonable to consider it as the current ‘default’ choice, in light of the wealth of available data, and in the interest of promoting comparability among studies.

## Methods

### Experimental animals

All experiments followed the standards of the German Federal Law on Care and Use of Laboratory Animals and were approved by the local government authorities (Lower Saxony State Office for Consumer Protection and Food Safety, approval number 33.19-42502-04-15/2042). A total number of 24 female adult Wistar rats (Charles Rivers Laboratories, Sulzfeld Germany) with a median body weight of 308 g (interquartile range 285-350 g) were used for this study. Rats were group-housed in cages with environmental enrichment, at a 12/12-hour light/dark cycle, with 20 – 24°C temperature and 45 – 55% humidity. Water and standard chow were provided ad libitum. The 24 rats were split into four equally sized groups, each assigned to a different protocol of medetomidine administration (see Table 1). No animal was excluded from the experiments or from the analysis. The investigators were not blind to the group allocation.

### Anesthesia and monitoring

Each animal was anesthetized on two sessions separated by a minimum of two weeks. The first session took place on a laboratory bench-top to accommodate unrestricted access to the animal and close monitoring of anesthesia duration and cardio-respiratory physiology (bench session). The functional imaging took place during the second session, performed inside a dedicated small animal MR system (fMRI session). All applied anesthetic protocols followed the same general outline, with isoflurane being used during preparation, and medetomidine during data acquisition (Fig. 1a).

Unconsciousness was induced in a chamber filled with 5% isoflurane and maintained throughout preparation with 2-3% isoflurane supplied through a nose cone. The eyes were covered with ophthalmic ointment to prevent them from drying. A cannula was inserted in the SC tissue of the left flank (protocol 1) or in a tail vein (protocols 2 – 4). Two subdermal needle electrodes were placed in the right forepaw, between the 2^nd^ and the 4^th^ digit. Monitoring equipment was attached, consisting of a rectal temperature probe, a pneumatic pressure sensor placed on the chest, and three SC needle electrodes for electrocardiogram (ECG). After fixing the aforementioned equipment with adhesive tape, the animal was transferred to a custom-built MRI-compatible rat bed.

Four different protocols of medetomidine administration were used: 1) SC with bolus; 2) IV with bolus; 3) IV no bolus; 4) IV lower dose. The detailed dosing for all protocols is given in Table 1. For protocols 1, 2, and 4, medetomidine (Dorbene vet, Zoetis Deutschland GmbH, Germany) was initially given as a bolus loading dose, followed by a gradual reduction of isoflurane and its eventual discontinuation 10 min later; continuous infusion of medetomidine commenced 15 min after the bolus. The bolus was omitted for protocol 3, with continuous infusion starting directly after the preparation phase, and isoflurane being gradually reduced to zero over the course of 20 – 25 min. Attempts to shut off isoflurane earlier led to fast breathing, indicative of an imminent wake-up (bench sessions). All doses were delivered using an MRI-compatible infusion pump (PHD 2000 Infuse/Withdraw; Harvard apparatus, Holliston, Massachusetts, USA). Since the start of medetomidine administration, heart rate (HR), respiratory rate (RR), and rectal temperature were monitored using the MR-compatible Model 1030 monitoring and gating system (Small Animal Instruments Inc., Stony Brook, NY 11790, USA). Rectal temperature was kept at 36.5 ± 1°C using a pad heated by circulating water. Anesthesia was maintained for six hours since the start of medetomidine administration, except for sessions in which the rat spontaneously woke up earlier. For bench sessions, such wake-ups were identified as spontaneous movements of the rat, and were found to be always preceded by the RR getting progressively faster and irregular. Since direct observation of minor animal movement is challenging during fMRI, the endpoint for fMRI sessions was set based on respiration (RR > 90/min, irregular, and continuously rising for at least 2 min). Anesthesia duration was defined as the time from the start of medetomidine administration (time = 0) till one of the aforementioned endpoints: spontaneous movement, rapid rise in RR, or passage of six hours. Upon reaching an endpoint, the animal was provided with 2% isoflurane through the nose cone and was disconnected from all electrodes, cannulas, and monitoring equipment. Finally, isoflurane was shut off and atipamezole (Atipazole, Prodivet pharmaceuticals, Belgium) was injected SC (0.25 mg/kg for protocols 1 – 3; 0.175 mg/kg for protocol 4) to facilitate a smooth wake-up.

The monitoring data (HR and RR traces) were recorded at a temporal resolution of 1 s and further processed with in-house python scripts as follows. Firstly, physiologically implausible values— corresponding to data acquisition errors—were dropped. Secondly, fMRI acquisition periods were removed from HR traces, since the rapidly switching magnetic gradients had introduced electrical noise in the ECG recording. Lastly, the traces were smoothed with an exponentially weighted moving average filter (smoothing factor α = 0.02). The processed HR and RR traces from bench and fMRI sessions closely resembled each other and were therefore pooled together. For each of the four medetomidine protocols the mean (± s.d.) HR/RR trace was calculated across all anesthesia sessions.

### MRI acquisition

The fMRI sessions were performed inside a 9.4 Tesla Bruker BioSpec MR system, equipped with the BGA12 gradient, and operated via Paravision 6.0.1 software. Signal was transmitted via a resonator volume coil (inner diameter 86 mm) and received by a rat brain 4-channel coil array (all equipment and software from Bruker Biospin MRI GmbH, Ettlingen, Germany). Approximately 10 min after the discontinuation of isoflurane, the rat was positioned in the isocenter of the MR system, with its body lying supinely and its head fixed with the help of a bite bar. After obtaining low-resolution images for object localization, a T2-weighted structural image was acquired using a TurboRARE sequence (repetition time 5.225 s, effective echo time 33 ms; 2 averages; RARE factor 8; 30 – 50 axial slices with a thickness of 0.5 mm; in-plane resolution 0.137 × 0.137 mm^2^; matrix size 256 × 256). A field map was measured and used for local shimming inside an ellipsoidal volume encompassing the rat brain (MAPSHIM). This was followed by multiple 330-second-long BOLD fMRI runs, repeated approximately every 10 min, until the experimental endpoint was reached. All fMRI runs were acquired with a single-shot gradient-echo echo planar imaging sequence (220 repetitions; repetition time 1.5 s; echo time 15 ms; flip angle 90°; 30 axial slices with a thickness of 0.5 mm, ascending interleaved slice order with no slice gap; in-plane resolution 0.2 × 0.2 mm^2^; matrix size 128 × 96; 4 dummy scans; bandwidth 375 kHz). The fMRI slices covered the entire rat brain, excluding the olfactory bulbs and the caudal 3/4 of the cerebellum. Consecutive fMRI runs were alternated between somatosensory fMRI with electrical forepaw stimulation (EFS-fMRI) and resting state fMRI (RS-fMRI) with no stimulus (Fig. 1b). This resulted in a total number of 283 EFS-fMRI and 295 RS-fMRI runs across all rats, spanning from 0.5 up to 6 hours since the start of medetomidine administration. EFS-fMRI runs included a baseline period of 60 s, followed by stimulation of the right forepaw in three 30 s blocks—each paired with 60 s of rest (Fig. 1c). Each stimulus block comprised square unipolar pulses with 3 mA amplitude and 0.3 ms pulse width, delivered at 9 Hz (Stimulus Generator 4002, Multi Channel Systems MCS GmbH, Reutlingen, Germany). The above parameters were chosen based on previous EFS studies in medetomidine-anesthetized rats^24,25,43^.

### MRI preprocessing

All MR images were first exported from Paravision to DICOM format and then converted to NIfTI (Neuroimaging Informatics Technology Initiative; http://nifti.nimh.nih.gov) using the dcm2nii (https://www.nitrc.org/projects/dcm2nii/) tool. The structural T2-weighted images were used to construct a study template with the help of the Advanced Normalization Tools software—ANTs (http://stnava.github.io/ANTs/). The structural image of one of the rats was chosen as the target reference space. Every other structural image was registered to the target in two steps: a linear rigid (3 translations and 3 rotations), and a non-linear symmetric diffeomorphic (SyN) registration^44^. All 24 structural images were averaged in the reference space to produce a mean anatomical image. A down-sampled (0.2 × 0.2 × 0.5 mm^3^) version of this image was used to create a brain mask and served as the study template to which all functional datasets were eventually registered. The fMRI image series were preprocessed using functions from multiple neuroimaging toolkits, combined into a pipeline with python’s Nipype library^45^. Images were corrected for slice timing with FSL (FMRIB Software Library, https://fsl.fmrib.ox.ac.uk/fsl/fslwiki/) and temporally filtered with AFNI (https://afni.nimh.nih.gov/). A high-pass filter of 0.01 Hz was used to remove slow temporal drifts, with an additional low-pass filter of 0.15 Hz being applied to RS-fMRI datasets only. Spatial smoothing (FSL) was performed using a 0.5 mm 3D Gaussian kernel. We chose to skip motion correction and regression of nuisance variables, as these preprocessing steps were shown to have little effect in anesthetized and head-fixed rodents^46^. A rigid transformation matrix was calculated between the mean image of each functional run and the native structural image. This matrix was combined with the previously calculated linear and non-linear transforms into a composite warp file, which was used to transform the preprocessed fMRI datasets into the study template space (ANTs).

### fMRI analysis and statistics

Areas activated during each EFS-fMRI run were identified via a first-level general linear model analysis carried out using FEAT (FMRI Expert Analysis Tool) Version 6.00, part of FSL^47^. The stimulus time course (1 during stimulation, 0 elsewhere) was convolved with a standard double-gamma hemodynamic response function to generate the model predictor. The resulting statistical maps were masked for brain, thresholded non-parametrically using clusters determined by z > 3.1 and a corrected cluster significance threshold of p = 0.05, binarized, and averaged across all 283 EFS-fMRI runs to construct an overall activation probability map. The region active in at least 30% of all EFS-fMRI runs (contralateral S1FL; see Fig. 3a) was taken as a region-of-interest (ROI). This ROI’s mean BOLD time course was extracted from each EFS-fMRI run, normalized to the pre-stimulus baseline, and averaged across the three stimulation blocks to produce an event-related average. The peak % signal change (peak ΔBOLD) was extracted from the event-related average, as a measure of BOLD response strength (Fig. 3b). We also performed a second-level fixed-effects analysis (FEAT) for each medetomidine protocol, by pooling the corresponding EFS-fMRI runs and computing the mean group effect. The resulting statistical maps were masked for brain and thresholded non-parametrically using GRF-theory-based maximum height thresholding with a (corrected) significance threshold of P=0.05.

Resting state functional connectivity (RSFC) analysis was performed using in-house python scripts. All 295 preprocessed RS-fMRI datasets were normalized by subtracting the temporal mean and dividing by the standard deviation. Normalized BOLD signal time courses were extracted from 28 ROIs (14 on each hemisphere), manually delineated based on the Paxinos-Watson rat brain atlas^48^ (see Fig. 5a, left). Pearson’s correlation coefficients were calculated for all ROI pairs, transformed into Fisher’s Z-scores, and stored as a pair-wise correlation matrix. The mean Z-score across all 378 unique ROI pairs served as a measure of global RSFC. To visualize the hierarchical structure of the functional connectome, each correlation matrix was also represented as a weighted graph, with the ROIs as nodes and the ROI pairs as edges. The edges were ranked by ascending correlation strength, and the rank was assigned as the numerical weight of the edge (strongest correlation to highest weight). Figure 5a (right) depicts a pair-wise correlation matrix and its graph representation for an example RS-fMRI run.

Statistical analysis was performed in R, version 3.2.5 (https://www.r-project.org/). The lme4 package^49^ was used to design a linear mixed effects model, separately for each of the four medetomidine protocols. This model can account for the repeated-measures design of the study and for missing data points (e.g. due to spontaneous earlier wake-ups). The EFS-evoked response strength (peak ΔBOLD) or the global RSFC (mean Z-score) were defined as response variables, time (onset of each fMRI run, relative to the start of medetomidine administration) as a fixed effect, and individual rat intercepts as random effects. The fit’s 95% confidence intervals were obtained through model-based bootstrapping, using the bootMer method of the lme4 package. To determine whether the response variables were significantly time-dependent, a Likelihood Ratio Test was used to compare each of the above models to a corresponding model lacking the time effect.

## Acknowledgements

This work was supported by the DFG (Deutsche Forschungsgemeinschaft) Research Center for Nanoscale Microscopy and Molecular Physiology of the Brain (CNMPB), and a DAAD (German Academic Exchange Service) stipend to N.S. We wish to thank Kristin Kötz, Kerstin Fuhrmann, and Luzia Hintz for their excellent technical assistance.

## Author contributions

The study was devised by J.B and S.B. Experiments and data analysis were performed by N.S. Results were discussed and interpreted by all co-authors. The manuscript was written by N.S, in consultation with S.B and J.B. All authors reviewed the manuscript.

## Competing interests

The authors declare no competing interests.

