## Supplementary for "Temporal stability of fMRI in medetomidine-anesthetized rats"

**Supplementary Table S1. (Dex)medetomidine administration protocols used by studies performing BOLD fMRI in rats.**

| Drug | Route* | Bolus dose (mg/kg) | Infusion rate (mg/kg/h) | Study |
|---|---|---|---|---|
| Medetomidine | SC | 0.05 | 0.1 | Weber et al., 2006[1] |
| | SC | 0.05 | 0.1 | Weber et al., 2008[2] |
| | SC | 0.05 | 0.1 | Zhao et al., 2008[3] |
| | IV | - | 0.1 | Pawela et al., 2008[4] |
| | IV | - | 0.1 / 0.3 / 0.1 to 0.3 | Pawela et al., 2009[5] |
| | SC | 0.05 | 0.1 | Seehafer et al., 2010[6] |
| | SC | 0.05 | 0.1 | Williams et al., 2010[7] |
| | SC | 0.05 | 0.1 | Angenstein et al., 2010[8] |
| | SC | 0.05 | 0.1 | Airaksinen et al., 2010[9] |
| | SC | 0.05 | 0.1 | Jonckers et al., 2011[10] |
| | SC | 0.5 | 0.1 | Majeed et al., 2011[11] |
| | SC | 0.05 | 0.1 | Airaksinen et al., 2012[12] |
| | SC | 0.05 | 0.1 | Krautwald and Angenstein, 2012[13] |
| | IV | 0.3 | - | Ciobanu et al., 2012[14] |
| | IP | 0.05 | 0.1 / 0.2 / 0.3 | Nasrallah et al., 2012[15] |
| | SC | 0.05 | 0.1 | Kalthoff et al., 2013[16] |
| | SC | 0.07 | 0.14 | Schwarz et al., 2013[17] |
| | SC | 0.05 | 0.1 | Angenstein et al., 2013[18] |
| | SC | 0.05 | 0.1 | Sekar et al., 2013[19] |
| | IV | 0.3 | - | Uhrig et al., 2014[20] |
| | IP | 0.05 | 0.1/ 0.2/ 0.3 | Nasrallah et al., 2014[21] |
| | SC | 0.2 | 0.1 | D'Souza et al., 2014[22] |
| | SC to IV | 0.05 | 0.1 | Duffy et al., 2015[23] |
| | SC | 0.05 | 0.1 | Sierakowiak et al., 2015[24] |
| | IP | 0.05 | 0.1 | Nasrallah et al., 2016[25] |
| | IV | - | 0.1 | Paasonen et al., 2016[26] |
| | SC | 0.05 | 0.1 | Scherf and Angenstein, 2017[27] |
| | SC | 0.04 | 0.05 | Albers et al., 2018[28] |
| | IV | 0.05 | 0.1 | Wang et al., 2018[29] |
| | IV | - | 0.1 | Shatillo et al., 2018[30] |
| | IV | - | 0.1 | Paasonen et al., 2018[31] |
| Dexmedetomidine** | IV | 0.05 | 0.05 | Fukuda et al., 2013[32] |
| | SC | 0.025 | 0.05 to 0.15 | Pan et al., 2013[33] |
| | SC | 0.025 | 0.05 | Chao et al., 2014[34] |
| | SC | 0.025 | 0.05 to 0.15 | Magnuson et al., 2014a[35] |
| | SC | 0.025 | 0.05 to 0.15 | Magnuson et al., 2014b[36] |
| | SC | 0.05 | 0.1 | Li et al., 2014[37] |
| | SC | 0.05 | 0.1 | Thompson et al., 2014a[38] |
| | SC | 0.025 | 0.05 | Thompson et al., 2014b[39] |
| | SC | 0.025 | 0.05 to 0.15 | Medda et al., 2016[40] |

* SC: subcutaneous; IV: intravenous; IP: intraperitoneal.

**Dexmedetomidine is the active ingredient of medetomidine, and has twice its potency; it is typically used at half the dosage

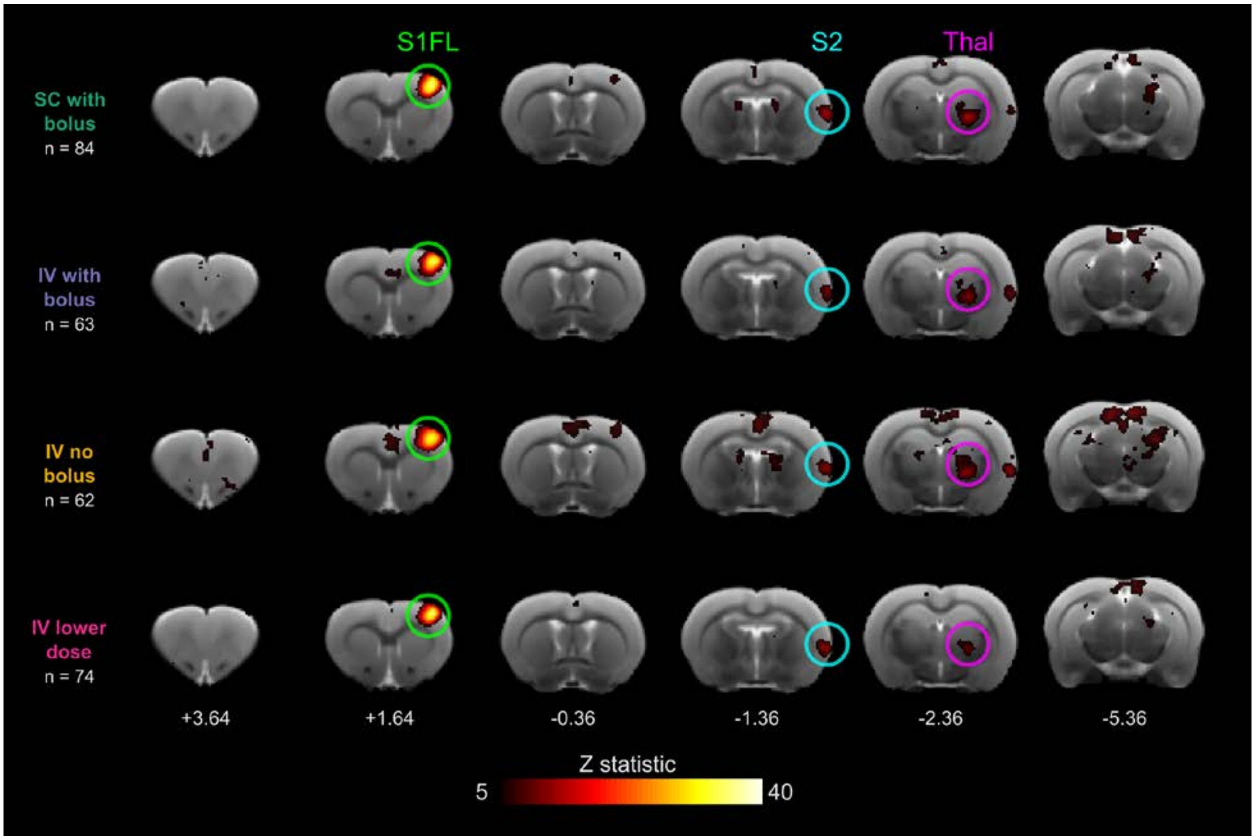


**Supplementary Figure S2. Areas activated by electrical forepaw stimulation (EFS): second level group analysis**

For each medetomidine protocol the corresponding first-level fMRI analysis results were pooled into a second-level fixed-effects analysis to compute the mean group effect (FMRI Expert Analysis Tool, version 6.00, part of FSL). The resulting statistical maps were masked for brain and thresholded non-parametrically using GRF-theory-based maximum height thresholding with a (corrected) significance threshold of P=0.05. The thresholded maps are shown overlaid on a T2-weighted structural study template. The approximate rostral-caudal position of each slice is given in mm relative to the bregma (based on the Paxinos-Watson rat brain atlas). Strong activation in the forelimb region of the somatosensory cortex (S1FL) is found in all protocols, with weaker activations in the secondary somatosensory cortex (S2), and the thalamus (Thal). Active clusters can also be found in cingulate and retrosplenial cortices, and in the brainstem.

### **References** (for Supplementary Table S1)

32. Fukuda, M., Vazquez, A. L., Zong, X. & Kim, S.-G. G. Effects of the alpha(2)-adrenergic receptor agonist dexmedetomidine on neural, vascular and BOLD fMRI responses in the

somatosensory cortex. *Eur. J. Neurosci.* **37**, 80–95 (2013).
